# Natural Bacteriocins as Potential Drug Candidates Targeting Core Proteins in Mastitis Pathogens of Dairy Cattle

**DOI:** 10.1101/2024.11.05.622021

**Authors:** Soharth Hasnat, Md. Morshedur Rahman, Farzana Yeasmin, Mohammad Jubair, Yosra A. Helmy, Tofazzal Islam, M. Nazmul Hoque

## Abstract

Mastitis poses a major challenge in the dairy industry, with rising antibiotic-resistant strains underscoring the urgent need for alternative antimicrobial strategies. This study aimed to (i) identify essential core proteins in clinical mastitis (CM)-causing pathogens using genomic approach, and (ii) assess the efficacy of natural antimicrobial peptides as novel therapeutic agents targeting the selected core proteins for the rational management of mastitis in dairy cows. Through a core genomic analysis of 16 CM-causing pathogens, including strains of *Staphylococcus aureus, S. warneri*, *Streptococcus agalactiae*, *S. uberis*, *Escherichia coli*, *Klebsiella pneumoniae*, *Pseudomonas aeruginosa*, *P. putida*, and *P. asiatica*, we identified 65 core proteins shared among these pathogens. Among them, ten proteins including PhoH, TrpB, FtsZ, HslV, HupB, RibH, InfA, MurA, GlxK, and Rho were found to be essential for the survival and virulence of these pathogens. Importantly, further novelty, resistance, and virulence assessments identified Rho and HupB as potential therapeutic targets. A comprehensive screening of 70 bacteriocin peptides (BPs) revealed 14 BPs that effectively interacted with both Rho and HupB proteins. Further analysis showed that BP8 and BP32 disrupt Rho protein function by blocking transcription termination process, while BP8, BP39, and BP40 prevent HupB from binding to DNA. These findings confirm the promising stability and efficacy of BP8 against both target proteins in CM-pathogens, highlighting it as a promising broad-spectrum therapeutic agent. Our computational study identified Rho and HupB as key proteins in CM-causing pathogens, which can be targeted by natural bacteriocins like BP8, suggesting its potential for developing effective and sustainable therapeutics against mastitis in dairy cattle.

**Author Summary:** Mastitis poses a significant threat to the global dairy industry, with rising antibiotic resistance necessitating alternative therapeutic strategies. This study identified essential core proteins in clinical mastitis-causing pathogens through a genomic approach and evaluated natural antimicrobial peptides (bacteriocins) as novel therapeutic agents. Through a core-genomic analysis, Rho and HupB were identified as key therapeutic targets. Bacteriocin peptides such as BP8 demonstrated promising efficacy by disrupting regular transcription termination process and DNA replication, offering a promising solution for next-generation mastitis therapies. The findings underscore the potential of BP8 as a sustainable, broad-spectrum antimicrobial agent, contributing to the rational management of mastitis in dairy cattle.

## Introduction

Bovine mastitis, a widespread inflammatory disease of the mammary gland, significantly impacts the dairy industry worldwide by reducing milk yield and quality, escalating treatment costs, and increasing culling rates (1, 2). This disease imposes a substantial economic burden on the dairy industry, with an annual loss between $19.7 billion and $32 billion USD globally (3), and an average cost of $147 per cow annually (4). Mastitis in dairy animals is a complex, multi-factorial disease caused by interactions between host, pathogen, environment, and gut microbiome dysbiosis, with bacterial pathogens being the primary contributors to its manifestations (5, 6). A plethora of microorganisms, including multidrug-resistant, pathogenic, opportunistic, emerging, and zoonotic bacteria, are the primary causal agents of mastitis in lactating mammals (7–9). Predominantly identified mastitis-causing bacterial pathogens include *Staphylococcus, Escherichia, Klebsiella*, *Streptococcus*, *Pseudomonas* etc., whose infections severely impact milk quantity, quality, and animal health (8, 10–12). The current treatment strategy for mastitis relies heavily on antibiotics, but the overuse and misuse of these drugs have significantly contributed to the rise of antimicrobial resistance (AMR) strains (13). The detrimental impact of antibiotics on the broader ecosystem has been well-documented in various studies (14–16). Moreover, the involvement of multiple pathogens in bovine mastitis, significantly complicates the infection process, as the increasing rate of antibiotic resistance diminishes the effectiveness of traditional antibiotics (17, 18). The rise of multidrug-resistant (MDR) strains highlights the looming ineffectiveness of antibiotics against mastitis pathogens, a situation exacerbated by decades of improper antimicrobial use in veterinary medicine (19). As a result, there is a global imperative to develop antibiotic alternatives for the treatment of mastitis, though significant progress remains elusive (20).

Given the diverse pathogens causing mastitis, there is an urgent need to modify traditional treatments to effectively combat MDR bacterial pathogens (8, 21). Identifying core proteins shared by the mastitis-causing pathogens, as core genomics approach is a promising strategy and has successfully identified therapeutic targets in previous studies (22, 23). The wide range of bacterial strains associated with mammary gland infections in cows (8, 10–12) has made it challenging to identify the core proteins shared by these mastitis-causing pathogens and thus yet remained unidentified. These core proteins are crucial for bacterial survival, and virulence, are essential for key metabolic pathways, and do not exhibit traits linked to AMR (24–28). For example, the transcription termination factor Rho is essential for factor-dependent transcription termination, a critical process that regulates gene expression in bacteria. Its role in coordinating the termination of RNA synthesis is fundamental to maintaining the balance of gene expression necessary for bacterial growth, adaptation, and overall cellular function (29–32). On the other hand, the nucleoid-associated protein (NAP) HupB plays an important role in maintenance of chromosomal architecture and in global regulation of DNA transactions in bacteria (33–35). Thus, identifying these core proteins as the therapeutic targets could revolutionize treatment strategies, providing a novel approach to effectively target the complex array of pathogens causing mastitis in dairy animals and other lactating mammals.

Natural antimicrobials like bacteriocins are emerging as promising alternatives to traditional antibiotics due to their potent activity against pathogenic bacteria and their relative safety for use in food and biomedical applications (36, 37). Bacteriocins are small, ribosomally synthesized peptides or proteins produced by bacteria (38, 39), targeting closely related species or even specific strains. Their effectiveness, coupled with their natural origin, makes them attractive candidates for combating AMR pathogens and enhancing food preservation without impacting beneficial microbiota (40). As research advances, harnessing bacteriocin peptides (BPs) from diverse sources (41) holds great potential for developing sustainable antimicrobial strategies that address current challenges in veterinary medicine. Therefore, proteinaceous antimicrobials could offer a novel and promising approach to managing bovine mastitis, marking a new era in treatment options. The feasibility of using BPs against mastitis-causing pathogens can be explored through computational approaches. Bioinformatics has already proven instrumental in identifying therapeutic targets, and recent advancements in identifying core proteins across multiple microbes have made this process even more robust (42). Additionally, the progress in computational biology has further streamlined the identification of effective proteinaceous antimicrobials against certain therapeutic targets (43, 44). This study aimed to develop a novel strategy for treating bovine mastitis by moving beyond conventional antibiotics. We employed *in-silico* methods to identify essential core proteins, specifically Rho and HupB, in bacterial pathogens previously reported to cause clinical mastitis in dairy animals (**S1 Table**). Additionally, we investigated natural antimicrobial peptides, such as bacteriocins (BPs), as potential therapeutic agents aimed at targeting these proteins. To achieve this, we analyzed the complete genomes of 16 pathogens responsible for clinical mastitis to identify their core proteins and effective bacteriocins. We then evaluated the potential of these bacteriocins against the Rho and HupB proteins, aiming to develop a more effective and sustainable strategy for the biorational management of mastitis in dairy cattle.

## Materials and methods

### Core genomic analysis

We selected the complete genomes of sixteen (N = 16) pathogenic bacterial strains associated with CM in dairy cows. These strains were isolated from the milk of CM-affected cows across eight different countries worldwide. The assembled genomes were obtained from the NCBI GenBank database, each associated with unique accession numbers (**S1 Table**). The annotated genomes were analyzed using the Pan-genome Explorer (PgE) as Prokaryotic genomes with gb extensions. To streamline the process and identify core proteins across these 16 genomes, we employed the PanACoTA pangenome software, applying a minimum percentage identity threshold of 50% (45, 46).

### Essential protein identification

Essential proteins are fundamental for cellular viability, representing the minimal set of genes necessary for the survival of a living cell (47). In our study, we focused on the essential proteins of the mastitis causing bacterial pathogens. We utilized the Database of Essential Genes (DEG) which catalogues 26,619 genes (48) deemed crucial for the survival of prokaryotic cells, including those associated with CM in lactating mammals. These essential genes are critical for understanding bacterial physiology and developing targeted antimicrobial strategies against CM pathogens. All essential protein sequences were retrieved from the DEG in FASTA format. To identify essential proteins in the mastitis-causing pathogens, we conducted a manual BLAST search via the Linux shell environment (49). The identified core proteins were used as query, while the essential protein served as the database. For conducting the manual BLAST analysis, we utilized BLAST+ executable v2.5.0, which was obtained from NCBI and run in the Linux shell environment (49). Subsequently, we created a customized database using the essential protein sequences. This allowed us to perform a BLASTp analysis of the core proteins from 16 CM-causing pathogens against the DEG. Essential proteins in these 16 pathogens were identified using an e-value threshold of 10⁻¹⁰. Additionally, a minimum cut-off score of 100 was applied in our analysis to further refine the identification of essential proteins (50).

### Enrichment analysis of the essential core proteins

ShinyGO v0.76 was utilized to gain insights into the molecular pathways and functions associated with the identified essential genes. This analysis offered a deeper understanding of the proteins involved in various biological roles, including interactions, reactions, metabolism, cellular processes, and disease mechanisms within the pathogens (51). The accession numbers of the identified essential proteins were obtained from NCBI and entered into ShinyGO, where a search was conducted for the best matching species (51). The program was executed using the recommended protein set as the background, with a false discovery rate (FDR) threshold of 0.05, and a pathway size range of 2 to 2000. The output generated included a relationship tree, protein-protein interaction network, KEGG pathway information, and Gene Ontology (GO) data for each protein. Essential proteins with available pathways and functional information were then selected for further analysis (52).

### Evaluation of novelty, antibiotic resistance, and virulence in identified essential proteins

To identify new therapeutic targets against bovine CM-pathogens, we avoided previously known targets listed in the DrugBank database (accessed on July 20, 2024), which contains over 14,000 proteins or drug targets (53). We conducted a manual BLASTp search comparing essential core proteins (with known metabolic functions) to therapeutic targets available in the DrugBank database (54). Novel therapeutic targets from the CM-causing pathogens underwent additional BLASTp analysis to assess potential antibiotic resistance features using proteins from the Comprehensive Antibiotic Resistance Database (CARD) (55). Furthermore, virulence proteins from the 16 CM-causing pathogens were identified by BLASTp against entries in the Virulence Factor Database (VFDB) (accessed on July 25, 2024) (56). Identified non-antibiotic resistance proteins (with known metabolic functions) were also queried against a custom VFDB-derived database. All BLASTp analyses utilized an e-value threshold of 10^-10^ and applied a minimum cut-off score of 100 for robust filtering of novel therapeutic proteins (57). This analysis identified two sets of essential core proteins (therapeutic targets), with each set comprising 16 proteins.

### Structural analysis of the final core proteins

After evaluation of novelty, antibiotic resistance, and virulence, structural preparation of the essential core proteins was performed through the web-based BLASTp suite (58). Core proteins from each set were uploaded as query sequences, and Protein Data Bank (PDB) database (59) was utilized to identify available structures. Among the final set of core proteins, experimentally confirmed PDB structures were only available for CM-causing *E. coli* and *S. aureus*. For the remaining core proteins of the each set, homology models were generated using the Rosetta program (60). Specifically, the experimentally validated structure <u>PDB ID 3ice</u> from *E. coli* was used as a reference for Rho core protein, while <u>PDB ID 8hd5</u> of *S. aureus* was used for HupB core protein. Structural comparison analysis was conducted using UCSF Chimera (61), where all protein structures within each core protein set were simultaneously input for comparison. To further explore structural and sequence variations, the multiple sequence alignment (MSA) module of BioLuminate v.5.5 was employed (62).

### Collection of bacteriocins and construction of the peptide library

Bacteriocin sequences (N = 32) were retrieved from the UniProt database (63) using advanced search options specifically designed to select bacteriocins, ensuring unbiased results. To identify the most effective regions, we fragmented the selected bacteriocins using PeptideMass (64). This tool processes protein sequences from the UniProt Knowledgebase (Swiss-Prot and TrEMBL) or user-submitted sequences by cleaving them with a selected enzyme and calculating the masses of the resulting peptides (64). We analyzed cysteine and methionine modifications by calculating peptide masses, treating cysteines as [M+H]^+^, and using monoisotopic mass calculations. Bacteriocins were cleaved with trypsin, allowing zero missed cleavages. The output displayed peptides (N = 70; BP1 – BP70) with masses greater than 500 Daltons, sorted either by peptide mass or by their sequence order within the protein. To generate three-dimensional (3D) structures, we used PEP-FOLD v3.5, a *de novo* peptide structure prediction tool (65). The raw amino acid (aa) sequences from our constructed bacteriocin library were input into the PEP-FOLD 3.5 server. The program was run with 200 ns simulations, and the models were sorted by sOPEP (66).

### Biochemical analysis of the BPs

A detailed biochemical analysis was conducted to characterize the physicochemical properties of each of the 70 BPs. For this purpose, we used the BIOCHEM peptide calculator (67), a specialized tool developed by Bachem, a leader in the production of high-quality active pharmaceutical ingredients (APIs) and pioneering chemical technologies. This tool was employed to determine the molecular weight, isoelectric point (IP), net charge, and average hydrophilicity of peptides within the BP library. Sequences were input using the one-letter code format with default N-terminal (H) and C-terminal (OH) configurations. Subsequently, PeptideRanker (66) was used to assess the bioactivity of each peptide. The PeptideRanker algorithm (68), which leverages the unique characteristics of active antimicrobial peptides, was crucial in predicting the antimicrobial potential of the identified BPs.

### Molecular docking interaction-based screening for protein-BP

Each peptide from the BP library was screened against the experimentally confirmed structures of Rho and HupB proteins, which represented each cluster of the CM-causing pathogens. This screening was conducted using Maestro v.14.0 from the Schrodinger suite (69), focusing on protein-BP molecular interactions. To investigate the interactions between the identified proteins (Rho and HupB) and the constructed BP library, the Rho and HupB proteins were designated as receptors, while the BPs were treated as ligands in the standard mode. Both receptors and ligands underwent a rapid structure preparation workflow to optimize them for molecular screening. The number of ligand rotations to be probed was set at 70,000, and the analysis was configured to return a maximum of 30 poses after screening. No specific attraction or repulsion restraints were applied during the screening process.

### Analysis of protein-BP interactions

To analyze the interactions between BP and the targeted proteins (Rho and HupB), we conducted a detailed chain-by-chain assessment. In each receptor-peptide interaction pose, the BPs were labelled as chain A, while the receptors were labelled as chain B (Rho) and chain C (HupB). We generated 30 distinct poses for each peptide-receptor interaction and meticulously examined them to evaluate interaction strength, focusing on hydrogen bonds (H-bonds), salt bridges, π-stacking, and Van der Waals clashes (70, 71). Advanced filtering techniques were applied to exclude poses with unfavourable clashes, retaining only those that exhibited H-bonds, π-π stacking, and salt bridges (70, 71). Finally, interaction diagrams were compiled for each of the energetically favourable poses.

### Analysis of the electrostatic potential surface for protein and BPs

The molecular electrostatic potential surface (EPS) was analyzed to visualize the three-dimensional (3D) charge distributions of interacting BPs and identified pathogenic core proteins (e.g., Rho and HupB). The EPS of the candidate BPs and their respective target proteins was computed using the Poisson-Boltzmann EPS structure analysis module in Maestro v.14 (69). This approach, based on the Poisson-Boltzmann equation (72), offered a structured insight into the electrostatic properties within these molecular systems.

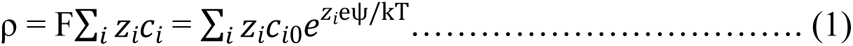

where: ρ (rho) is the charge density; F is Faraday’s constant; *i* is the ionic species; *Z*_*i*_ is the valence of ion species; *C*_*i*0_ is the reference concentration of ion species; e is the elementary charge; ψ (psi) is the electrostatic potential at a given point; k is the Boltzmann constant; T is the absolute temperature in Kelvin.

The EPS visualizations played a critical role in predicting the mechanisms underlying BP-protein interactions, providing valuable insights for optimizing the dynamics of peptide-protein interactions.

### Stability analysis of the protein-BP complexes

To evaluate the stability of the receptor (Rho and HupB)-BP complexes, we conducted molecular dynamics simulations (MDS) on the top protein-BP interaction poses. These simulations were performed using the Desmond module of Schrödinger (release 2024-2) on a Linux-based system (73). The MDS process began with protein preparation, during which any missing side chains were reconstructed. Overlapping hydrogens were automatically optimized, followed by restrained minimization to finalize the structure. Water molecules located more than 5 Å from the complexes were removed to streamline the simulation. The processed complexes were then prepared using the Desmond System Builder, with TIP3 set as the solvation model. An orthorhombic box with a 10 Å margin was constructed to simulate interaction strength. The system was built using the buffer calculation method, employing the OPLS4 force field, and neutralized by the addition of Na+ or Cl− ions (74). The constructed systems were subsequently loaded into the MDS workflow, with the simulation time set for 200 ns and 1000 frames (71, 75). During this analysis, the NPT ensemble method (constant number of particles, pressure, and temperature) was employed, maintaining a temperature of 300K and a pressure of 1 bar (75). The simulation was executed using an NVIDIA RTX A6000 GPU. At the conclusion of the MDS, the frames were collected and analyzed using a simulation interaction diagram to assess the strength and fluctuations of the peptide-receptor interactions (75).

### Feasibility assessment of BPs targeting the Rho protein

The Rho protein (PDB ID 3ICE) consists of six identical chains, with RNA bound at the centre of the hexamer. We isolated the A chain, along with its bound RNA and ADP using Maestro v14.0 (69), to evaluate the stability of nine interacting BPs that exhibited stable interaction with the Rho protein in initial screening and MDS. The RMSD values for the Rho-BP complexes, ranging between 1-5 Å, suggested adequate stability for further screening using BioLuminate v.5.5 (62). This screening involved evaluating 70,000 ligand rotations, generating up to 30 poses per ligand, without applying any specific attraction or repulsion constraints. A parallel screening was conducted on the Rho-RNA-ADP complex without any BPs. To assess the overall stability, a 200 ns MDS was performed using the Desmond engine in the Schrödinger suite (73). The Desmond SID module was subsequently employed to further assess the dynamic stability of the system.

## Results

### Sixty-five set of proteins identified in mastitis causing pathogens through core genomic approach

A comprehensive genomic analysis of 16 CM-causing pathogens (**S1 Table**) was performed using the PanExplorer (PgE) web-based tool (46). These pathogens include three strains each of *S. aureus*, *E. coli*, and *K. pneumoniae*, along with two strains of *S. agalactiae*, and single strains from *S. warneri*, *S. uberis*, *P. aeruginosa*, *P. putida*, and *P. asiatica* (**S1 Table**). These strains were isolated and sequenced from milk samples of cows suffered from CM across diverse geographical locations worldwide. This analysis generated a heatmap with hierarchical clustering, providing a comparative visualization of the genomic similarities and differences among the pathogens (**Fig. 1A**). The dendrogram generated by PgE grouped these strains into clusters according to their genetic similarity. The analysis of the complete genomes revealed that the core genomes of *S. aureus*, *S. warneri*, *S. agalactiae*, and *S. uberis* strains are genetically similar, as illustrated in the heatmap (**Fig. 1A**). Furthermore, our analysis revealed that *E. coli* strains exhibited closer genetic relationships with *P. aeruginosa*, *P. putida*, and *P. asiatica* strains than *K. pneumoniae* strains (**Fig. 1A**). On average, the gram-negative pathogens including strains of *K. pneumoniae*, *E. coli*, *P. putida*, *P. asiatica*, and *P. aeruginosa* had higher gene counts, with each strain containing over 4,000 genes, compared to gram-positive strains of *S. aureus*, *S. warneri*, *S. agalactiae*, and *S. uberis*, with fewer than 3,000 genes per strain. Among the 16 strains analyzed, *S. uberis* 2062 had the lowest gene count with 1,895 genes, while *P. aeruginosa* 2011C-S1 had the largest count, containing 6,192 genes (**Fig. 1B**). Core genome analysis identified only 65 core genes (proteins) shared across all the pathogens. Additionally, a significant number of genes were classified as dispensable (partially core), indicating that they were not consistently present across all 16 CM-causing pathogens (**Fig. 1B**).

**Fig. 1.**
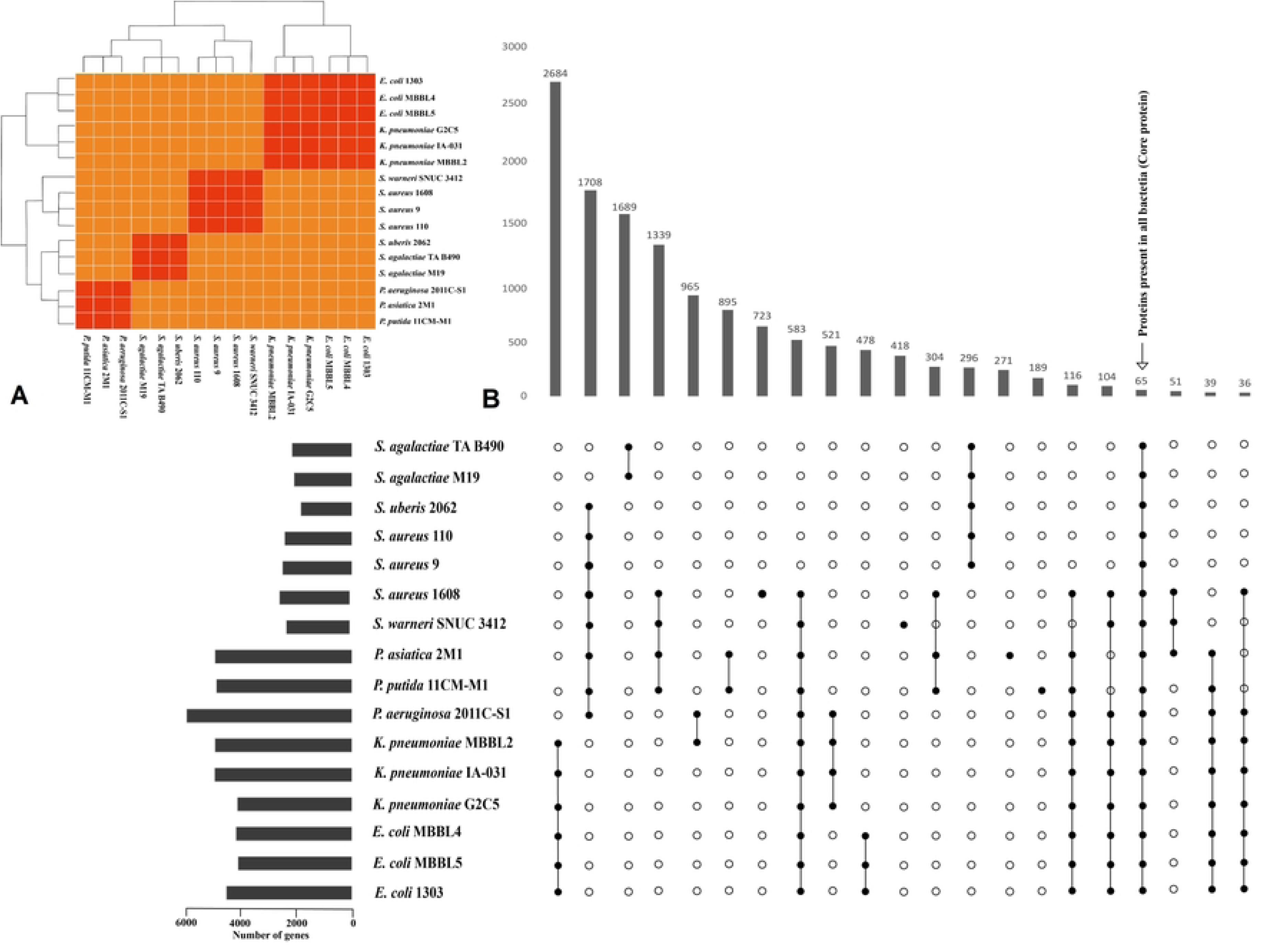
Shared genome and core proteins in 16 mastitis associate pathogens. (A) Clustered heatmap illustrating genomic similarities and highlighting shared regions (red boxes) among 16 pathogens, revealing their common genetic elements. (B) Upset plot comparing the distribution of common and core proteins among 16 pathogens. Vertical bars in the upper right panel showing the number of core proteins in 16 genomes (based on the presence and absence matrix). Box plots in the lower right panel represents the mean distribution of core proteins in the 16 CM-associated pathogens. Horizontal bars in the lower left panel provides a comparative overview of genomic contents.

### Ten essential core proteins identified in mastitis causing pathogens

We identified 65 sets of core proteins, each comprising 16 proteins, across the CM-causing pathogens. However, the essentiality of these proteins for bacterial survival was initially unknown. Upon aligning these core proteins with the database of prokaryotic essential proteins, it was determined that only ten of the 65 core proteins were essential for the survival of these pathogens. The ten essential core proteins identified were the PhoH family protein, tryptophan synthase subunit TrpB, cell division protein FtsZ, ATP-dependent protease subunit HslV, HU family DNA-binding protein HupB, ribityllumazine synthase RibH, translational initiation factor InfA, peptidoglycan biosynthesis enzyme MurA, glycerate kinase GlxK, and transcription termination factor Rho. Each type of core essential protein shared at least 60% similarity with corresponding proteins in the database of essential proteins (**S2 Table**). These strong similarities suggest that these core proteins are involved in critical metabolic pathways essential for the survival of the 16 CM-causing pathogens.

### Enrichment analysis elucidated the role of core proteins in gene expression of mastitis pathogens

All ten essential proteins (**S2 Table**) underwent enrichment analysis to reveal their roles in the cellular activities of CM-pathogens. We found that these proteins are involved in 18 cellular processes, suggesting that each protein may play multiple molecular functions (**Fig. 2A**). High fold enrichment values for replication, transcription, and translation processes indicate a significant involvement of the essential core proteins in these pathways. Remarkably, these proteins were strongly linked to translation initiation process (InfA), transcript initiation and termination (Rho), riboflavin biosynthesis (RibH), DNA binding and packaging (HupB), cell division (FtsZ), protein catabolic process (HsIV), and biosysnthesis (TrpB, MurA, GlxK, PhoH), among other molecular functions (**Fig. 2A**). The low FDR score (1.44) for the selected essential proteins confirms the accuracy of the enrichment analysis. To determine the specific molecular functions and assigned pathways of these essential genes, we examined KEGG pathway (KO) information and Gene Ontology (GO) data. We identified unique KO and GO IDs with significant e-values for each essential gene, highlighting their importance in pathogen survival (**Fig. 2B**)

**Fig. 2.**
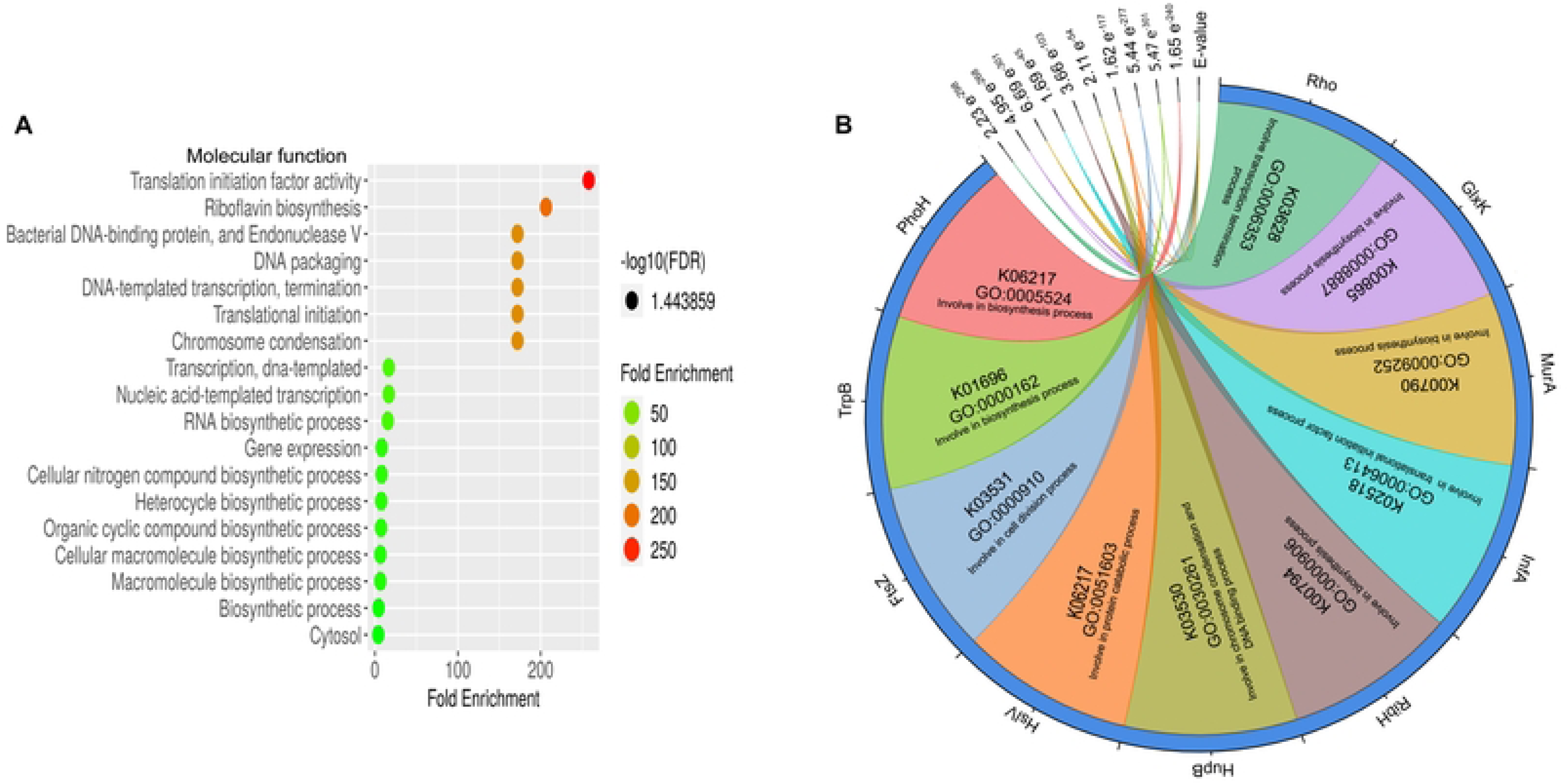
Analysis of molecular functions and pathways in core proteins. (A) Detailed molecular function analysis displaying fold enrichment for each essential gene, underscoring their functional roles and relative importance. (B) KEGG Orthology (KO) and Gene Ontology (GO) annotations for each essential gene, accompanied by significant e-values, indicating the statistical relevance of these associations.

### Novelty, resistance, and virulence assessments reveal two potential therapeutic targets for mastitis pathogens

We evaluated the novelty, antimicrobial resistance, and virulence of ten selected proteins using various computational tools. In the novelty analysis, we used ten essential proteins such as PhoH, TrpB, FtsZ, HslV, HupB, ribH, InfA, MurA, GlxK, and Rho (**Fig. 2B**) that did not correspond to any known therapeutic targets, based on established therapeutic targets available in <u>DrugBank database</u>. The findings of the novelty assessment suggested that these novel proteins could be potential therapeutic targets. However, since a viable therapeutic target must lack antibiotic resistance features, we compared these novel proteins with those in the ARGs database (e.g., CARD) and found that PhoH, MurA, and GlxK have over 50% sequence similarity with known antibiotic resistance proteins. As a result, these three were excluded from consideration. Given that mastitis is an inflammatory disease, we focused on proteins with virulent properties as potential therapeutic targets. Among the remaining seven proteins, HU (histone-like) family DNA-binding protein HupB and the transcription termination factor Rho, exhibited substantial virulence features. Thus, these two core proteins (Rho and HupB) were identified as novel therapeutic targets against CM-pathogens.

### Molecular architecture of core proteins reveal functional conserveness

Two core protein sets were ultimately identified, each comprising 16 proteins, one set consisting of Rho proteins and the other of HupB proteins. Superimposing the structures of 16 Rho proteins revealed five distinct structural isoforms across the CM-pathogens, represented by colors gold, green, pink, brick red, and blue (**Fig. 3A**). Notably, *S. aureus*, *S. warneri*, *S. agalactiae*, and *S. uberis*, despite being different species, shared identical Rho protein. Conversely, the Rho protein of *P. asiatica* 2M1 showed the most notable structural divergence compared to other Rho isoforms (**Fig. 3A**). The remaining Rho proteins exhibited greater structural similarity, except for the Rho proteins from *E. coli* and *P. putida* (**Fig. 3A**). The subsequent MSA analysis provided detailed insights into the amino acid (aa) sequence similarities and variations among the 16 Rho proteins. The Rho protein from *P. asiatica* 2M1 displayed only 51% aa sequence similarity compared to the others. In contrast, three *E. coli* strains and one *P. putida* strain shared a 53% similarity in their aa sequences, while three *K. pneumoniae* strains and one *P. aeruginosa* strain showed a higher similarity range of 80–81%. Remarkably, three *S. aureus* strains, two *S. agalactiae* strains, and single strains of *S. warneri* and *S. uberis* exhibited complete identity, with 100% sequence similarity in their Rho protein structures (**Fig. 3B**). These MSA results confirmed the structural similarities and discrepancies among the Rho proteins of the selected CM-causing pathogens (**Figs. 3A-B**). Further molecular studies focused on an experimentally confirmed Rho protein from *E. coli* (<u>PDB ID 3ice</u>), which showed 100% aa sequence similarity with the Rho proteins of CM-causing *E. coli* (**Fig. 3C**). Similarly, the 16 HupB proteins were superimposed, revealing six distinct structural isoforms (**Fig. 3D**). The structures of HupB proteins from different strains of *S. aureus*, *S. warneri*, *S. agalactiae*, and *S. uberis* were found to be completely identical. On the other hand, the HupB protein of *P. putida* 11CM-M1 showed the most notable structural divergence compared to other HupB isoforms (**Figs. 3D**). Further MSA analysis revealed aa sequence similarities and differences, showing that various strains of *S. aureus*, *S. warneri*, *S. agalactiae*, and *S. uberis* had completely identical sequences (100% similarity). In contrast, the HupB proteins of other strains shared approximately 55–80% aa sequence similarity, except for the HupB protein from *P. putida* 11CM-M1, which displayed only 50% sequence identity (**Fig. 3E**). An experimentally confirmed HupB protein from *S. aureus* (<u>PDB ID 8hd5</u>) was found to share 100% aa sequence similarity with the HupB proteins from the CM-causing gram-positive pathogens of this study (**Figs. 3E-F**). Therefore, PDB ID 8hd5 was selected for further molecular studies.

**Fig. 3.**
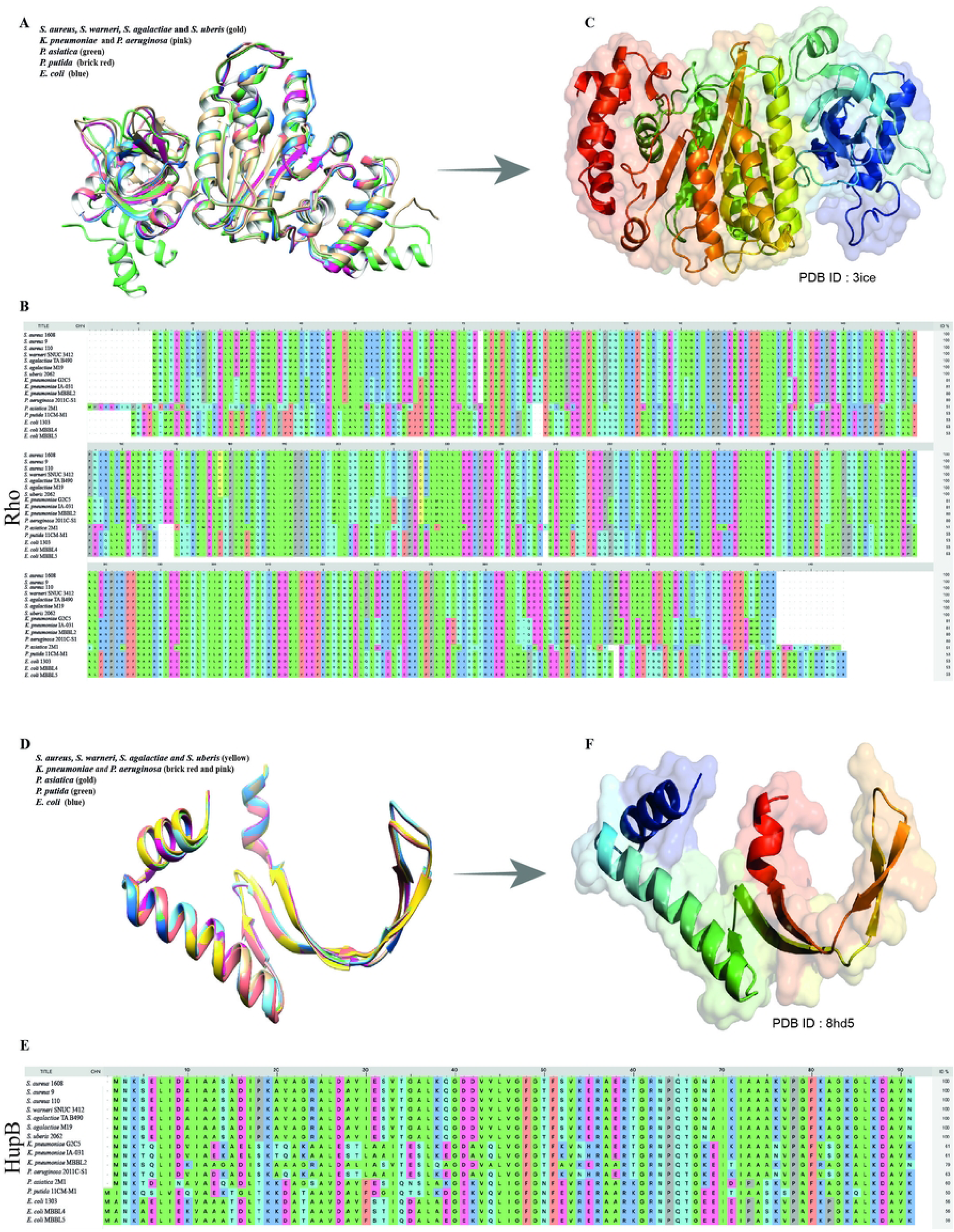
Structural comparison and multiple sequence alignment (MSA) of 16 Rho and HupB proteins, illustrating their diversity, sequence similarities, and highlighting isoforms and conservation. (A) Structural comparison of the 16 Rho proteins reveals five distinct structural isoforms. (B) Graphical representation of MSA for Rho proteins, indicating percentage identities to highlight similarities and differences among the 16 proteins. (C) Structural comparison of the 16 HupB proteins identifies five distinct structural isoforms. (D) MSA of HupB proteins with percentage identities, demonstrating the degree of sequence conservation among the 16 proteins.

### Biochemical characteristics of bacteriocin peptides

Understanding the biochemical properties of bacteriocin peptides (BPs) is crucial for developing and applying them as potential therapeutic agents. Our comprehensive extraction method identified 70 BPs from 32 bacteriocins. Each BP underwent thorough biochemical characterization, revealing amino acid (aa) counts ranging from 5 in the smallest peptide to 29 in the largest **(S3 Table**). The molecular weight of these BPs varied significantly, with BP47 being the heaviest at 3115.64 g/mol and BP61 the lightest at 545.64 g/mol. Detailed molecular weight data for all 70 BPs is available in **S3 Table**. Isoelectric point (IP) measurements, a key biochemical attribute, revealed that 29 of the BPs had an IP greater than 7, while the others had values below this threshold. BP19 exhibited the highest isoelectric point (IP) at 12.49, while BP69 recorded the lowest. The BPs were further classified based on their net charge: 29 were negatively charged (ranging from -0.05 to -2.00), 11 were neutral (0.0), and 30 were positively charged (ranging from 0.05 to 2.00) (**S3 Table**). Hydrophilicity, a critical factor in antimicrobial activity, also showed considerable variations. Among the identified BPs, 45 exhibited negative hydrophilicity values (-0.03 to -1.72), three were neutral (0.00), and 22 had positive values (0.09 to 1.76). The proportion of hydrophilic aa relative to the total number of aa for each BP is also shown in **S3 Table**. The highest ratio of hydrophilic aa was found to be 88% for BP48, while the rest had lower ratios. Notably, two BPs (BP9 and BP60) had no hydrophilic amino acids at all. In the bioactivity assessment, BP8 (score 0.99 out of 1) was identified as the most bioactive antimicrobial peptide, while BP17 (score 0.03 out of 1) was the least active among the 70 BPs analyzed (**S3 Table**).

### Nine BPs reveal robust interactions with Rho protein in molecular screening

The molecular screening of the BPs against the Rho protein revealed that nine BPs namely BP2, BP8, BP26, BP30, BP32, BP35, BP41, BP54, and BP64, effectively interacted and formed potential inhibitory complexes. BP2 formed a complex with the Rho protein through specific molecular interactions involving its aa residues TYR1, TYR2, ASN4, VAL6, and LYS10, which formed key contacts with the GLN59, ARG109, and ALA74 residues of the Rho protein. These interactions suggest a critical role for these residues in mediating the BP2-Rho protein complex formation. Notably, the side chain atoms of ARG109 in the Rho protein formed three hydrogen bonds (H-bonds) with TYR2, ASN4, and VAL6 of BP2. Additionally, the backbone atoms of TYR1 and LYS10 established H-bonds with GLN59 and ALA74, respectively, in the BP2-Rho complex (**Fig. 4A**). The most robust interaction was observed in the BP8-Rho complex, where eight aa of the Rho protein formed side chain-associated H-bond interactions with eight corresponding aa of BP8. These interactions were established between the following pairs: ASN4--GLU5, TYR5--SER50, GLY6--ASP52, GLY8--LYS249, SER12--GLN220, TRP15--GLN8, ASN20--ARG224, and LYS258--ASN26 (**Fig. 4B**). BP26 also exhibited interactions with the Rho protein, where the backbone atoms of VAL7 and ARG367 formed H-bonds, while the association of LYS9, ARG367, and GLU369 was mediated by salt bridges (**Fig. 4C**). BP30 formed seven H-bonds with the Rho protein through the side chain atoms of the following aa pairs: GLU1--GLU368, ASN2--GLU342, ALA4--LYS335, ILE6--HIS338, and MET18--TYR332. Additionally, a single π-π stacking interaction was identified between TRP11 of BP30 and HIS338 of the Rho protein (**Fig. 4D**). BP32 exhibited a complex array of four different types of chemical interactions with the Rho protein. Specifically, the side chain atoms of CYS1 and VAL3 formed H-bonds with GLU368 and LYS385, respectively. Moreover, the backbone atoms of CYS1 formed an H-bond with LEU164, while ALA8 established an H-bond with GLU369. LYS9 displayed coupled interactions, primarily consisting of salt bridges and H-bonds, with ARG367 and ASP374. Additionally, TRP5 engaged in a π-π stacking interaction with TRP381 of the Rho protein (**Fig. 5E**). BP35 demonstrated binding with the Rho protein, where the side chain of GLU3 formed a hydrogen bond (H-bond) with TYR80, and the backbone atoms of THR5 formed an H-bond with ARG109 (**Fig. 5F**). In the BP35-Rho complex, ARG109 of the Rho protein exhibited a coupled interaction (CI), involving both an H-bond and a salt bridge with GLU2 of BP35. The BP41-Rho complex involved interactions with 11 aa. Specifically, GLY4 interacted with ARG221 and ASP233 interacted with ALA12 and THR23 through H-bonds formed by side chain atoms. Additionally, SER21 interacted with ASP69 and TYR interacted with GLY225 through H-bonds involving backbone atoms. A single CI was observed between GLU17 and ARG238 within the BP41-Rho complex (**Fig. 4G**). A similar interaction pattern was observed in the BP54-Rho complex. Here, the backbone atom of MET1 formed a H-bond with GLY107, while the side chain atoms of LYS6 established an H-bond with GLN59. LYS6 was also involved in a CI with ASP60 (**Fig. 4H**). The BP54-Rho complex was stabilized by interactions involving seven aa. Notably, PHE36 and PHE111 of BP54 interacted with TYR6 of the Rho protein through a π-π stacking bond. Additionally, the side chain atoms of LEU9 formed an H-bond with LYS40, and GLU106 engaged in a salt bridge interaction with LYS11 (**Fig. 4I**). The biochemical and bioactivity characteristics of the nine interacting BPs were also studied individually to understand their effectiveness against the Rho protein. The sequence length, molecular weight, bioactivity score, isoelectric point and net charge of these BPs ranged from 5 to 27 nucleotides, 589.76 to 2936.28 g/mol, 0.056 to 0.999, 4.3 to 10.45, -1 to 2.05, and -0.58 to 1.46 (**Table 1**).

**Fig. 4.**
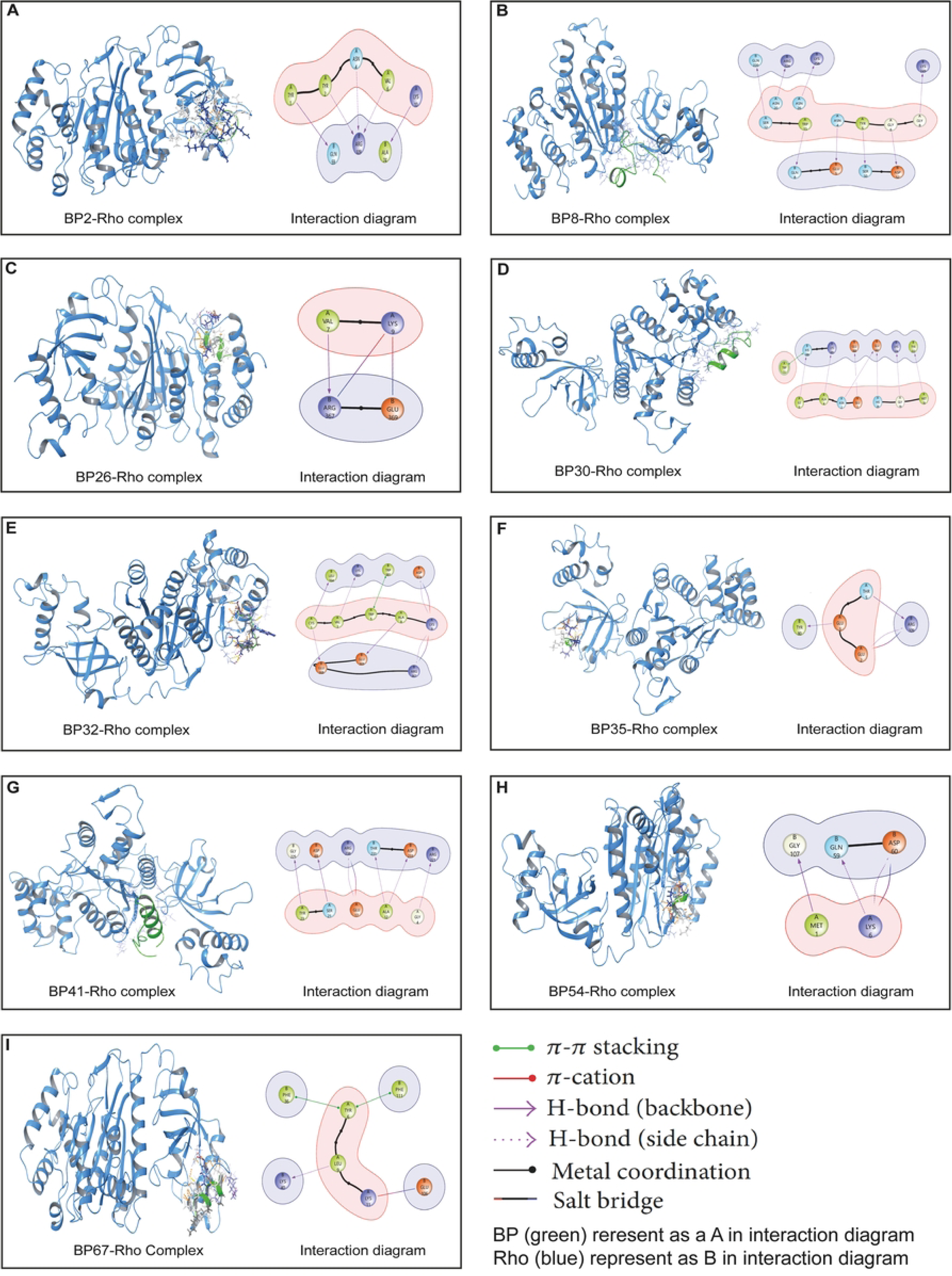
The molecular interactions between nine bacteriocin peptides (BPs) and the Rho protein. (A) BP2-Rho complex, (B) BP8-Rho complex, (C) BP26-Rho complex, (D) BP30-Rho complex, (E) BP32-Rho complex, (F) BP35-Rho complex, (G) BP41-Rho complex, (H) BP54-Rho complex and (I) BP67-Rho complex, with interaction diagrams illustrating the binding details.

**Fig. 5.**
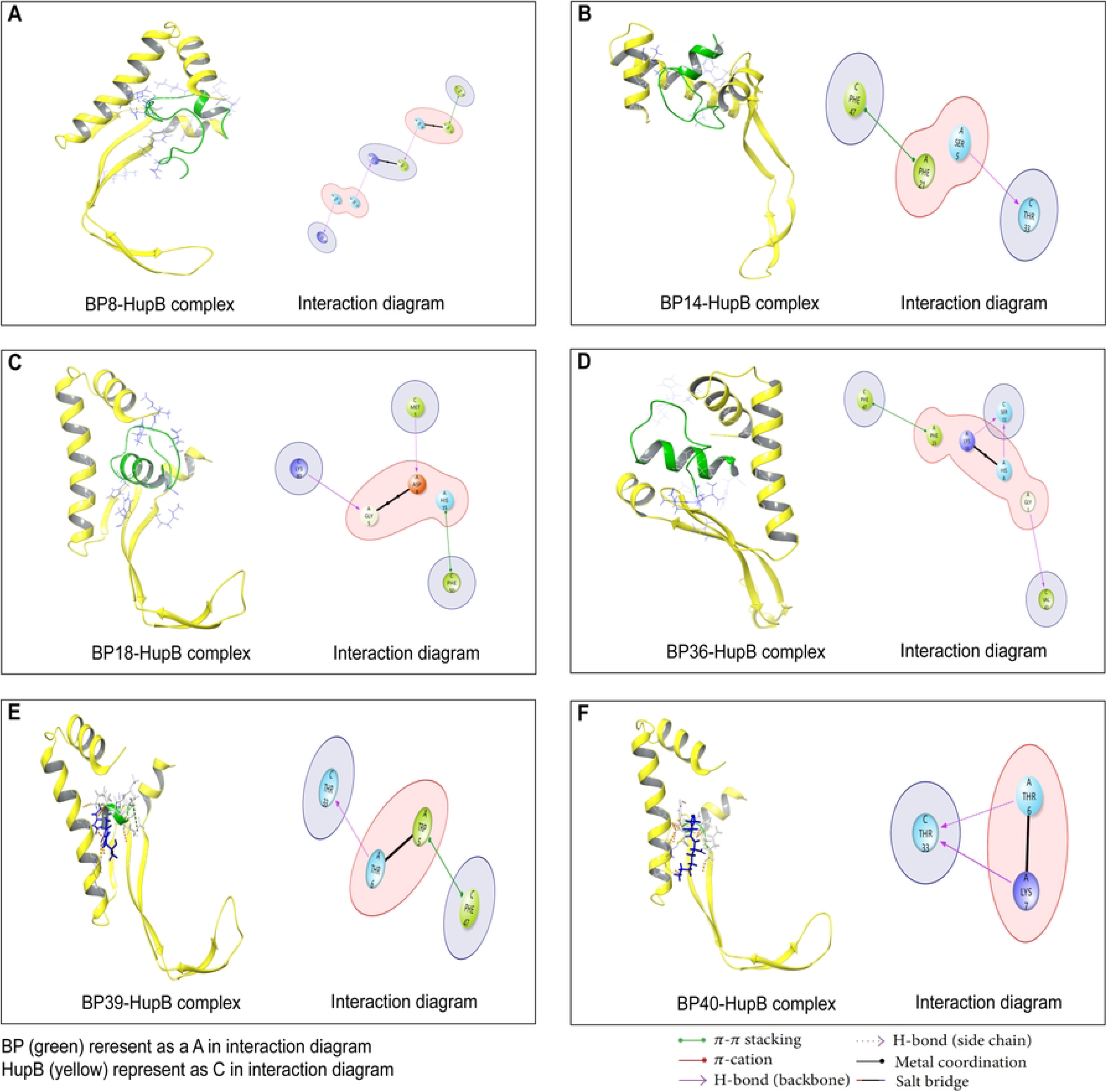
The molecular interactions between six bacteriocin peptides (BPs) and the HupB protein. (A) BP8-HupB complex, (B) BP14-HupB complex, (C) BP18-HupB complex, (D) BP36-HupB complex, (E) BP39-HupB complex, and (F) BP40-HupB complex, with interaction diagrams illustrating the binding details.

**Table 1.**
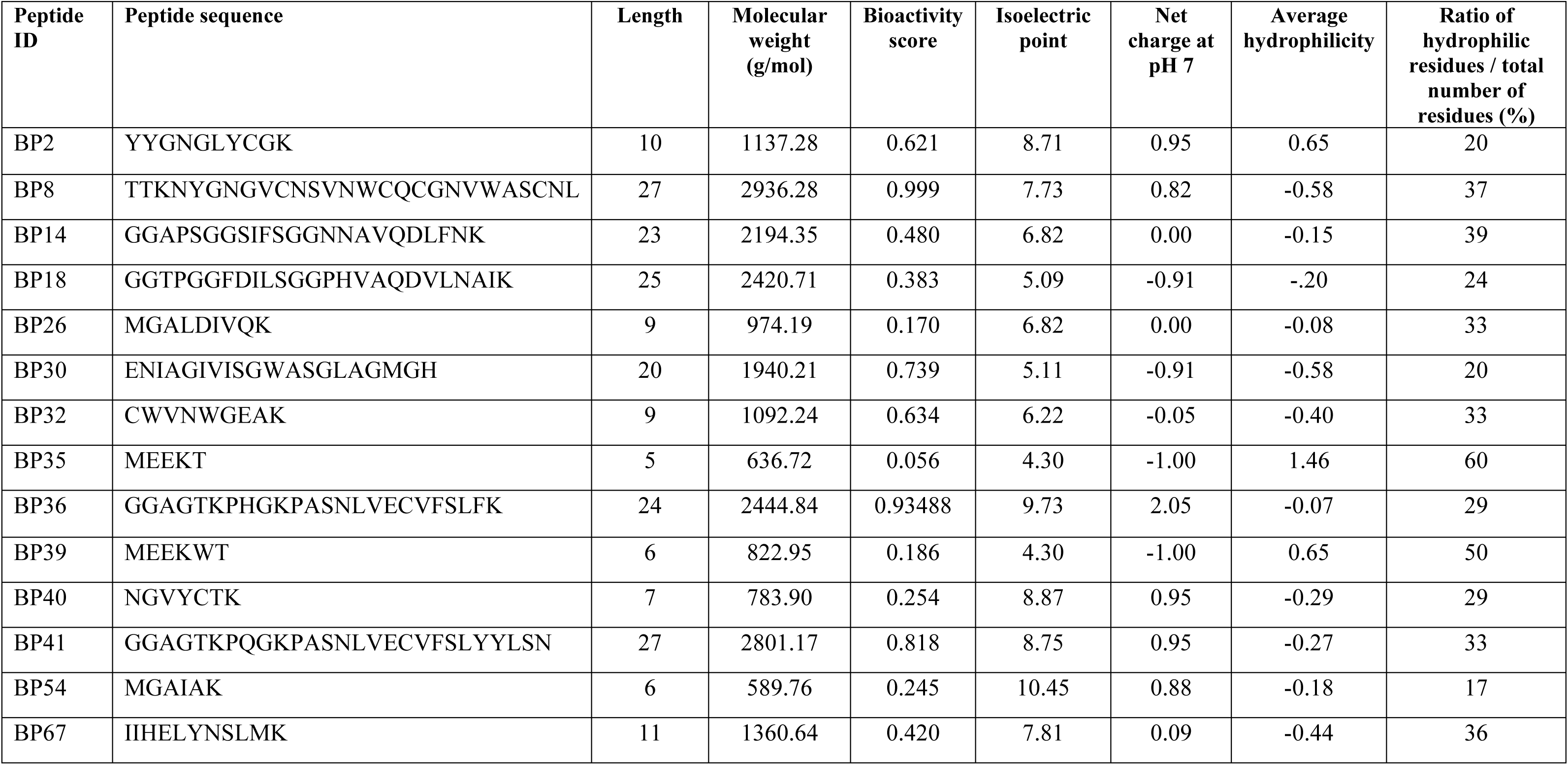
Biochemical properties of bacteriocin peptides (BPs) interacting with Rho and HupB proteins.

### Six BPs reveal robust interactions with HupB protein in molecular screening

The 70 identified BPs were also screened against the HupB protein, revealing interactions between six BPs and the HupB protein. BP8 was found to form H-bonds with ASN4--LYS53, SER12--LYS3, and SER24--MET1 residues of the HupB protein via side chain atoms. Additionally, a π-π stacking interaction was observed in the BP8-HupB complex between the TRP22 and PHE79 residues (**Fig. 5A**). In the BP14-HupB complex, only two interactions were identified: an H-bond mediated by the side chain of SER5 with THR33 and a π-π stacking interaction between PHE21 and PHE47 (**Fig. 5B**). The BP18-HupB complex was stabilized by interactions involving six aa. The backbone atoms of LYS80 formed an H-bond with GLY5, while the side chain of MET1 formed an H-bond with ASP8. Additionally, HIS15 of BP18 engaged in a π-π stacking interaction with PHE50 of the HupB protein (**Fig. 5C**). In the BP36-HupB complex, three aa of BP36 formed side chain-associated H-bond interactions with two corresponding aa of HupB. These interactions occurred between GLY1--VAL43, HIS8--SER13, and HIS8--LYS10. A single π-π stacking interaction was also identified between PHE23 and PHE47 (**Fig. 5D**). The BP39-HupB complex involved only four aa, where the side chain of THR6 interacted with THR33 via H-bonding, and TRP5 formed a π-π stacking interaction with PHE47 (**Fig. 5E**). In the last BP40-HupB complex, the aa THR6 and LYS7 of BP40 interacted with THR33 of the HupB protein only through H-bonding (**Fig. 5F**). The biochemical and bioactivity properties of these six interacting BPs were also individually analyzed to identify the factors contributing to their effectiveness against the HupB protein (**Table 1**).

### EPS analysis unveils interacting polar surfaces suitable for better protein-peptide interactions

All 14 interacting BPs, along with the core proteins Rho and HupB, were subjected to EPS analysis to assess their three-dimensional charge distributions. The EPS analysis revealed insights into the relative polarity of the molecules during their interactions (**Fig. 6**). For example, BP2 exhibited EPS values ranging from -27.09 to 27.09 a.u., with red regions indicating electronegative atoms (primarily oxygen and nitrogen) corresponding to negative EPS values. In contrast, white regions denoted neutral surface areas, while blue regions represented areas with positive potentials. BP8, which demonstrated the highest interaction with the Rho protein, displayed more extensive nucleophilic site (red regions) across its surface compared to the other BPs interacting with Rho. BP8 exhibited EPS values ranging from -32.45 to 37.7 a.u. BP26, characterized by large electrophilic site (blue regions), had EPS values between -35.07 and 23.22 a.u. BP30, with a predominantly negative surface, ranged from -20.47 to 23.5 a.u. BP32 and BP35, both largely covered by blue regions, had EPS values of -30.23 to 28.4 a.u. and -27.58 to 19.23 a.u., respectively. BP41 was mainly dominated by negative charges, with values from -30.65 to 31.2. BP54, showing prominent red regions, ranged from -15.14 to 20.11 a.u. Finally, BP67, mostly covered by positive charges with some dark red areas, exhibited EPS values from -36.85 to 25.77 a.u. (**Fig. 6A**). The EPS values for both the Rho and HupB proteins were also calculated. The Rho protein exhibited a range of EPS values from -60.11 to 48.37 a.u., displaying a mixed surface distribution. Several small regions were marked with dark red, indicating strongly negative charges, while the rest of the surface was covered by a slightly blue hue, representing positive charges. Likewise, the surface of the HupB was mainly moderate blue with a few dark red spots with EPS values ranging from -41.64 to 35.94 a.u., indicating uniform charge distribution (**Fig. 6B**). We further examined the EPS of various BPs in their interactions with HupB protein. BP8 interacted with both proteins without significant EPS changes for HupB. BP14 showed EPS values from -41.71 to 27.26 a.u., mostly dark blue, indicating strong positive potential. BP18 had the most negative EPS value at -54.03 a.u., with a light blue to white surface and a few dark red spots. BP36 and BP39 exhibited mixed charges, while BP40 showed negative charge dominance with EPS values from -22.38 to 27.5 a.u. (**Fig. 6C**). Overall, BP8 exhibited the highest EPS values among all the interacting BPs.

**Fig. 6.**
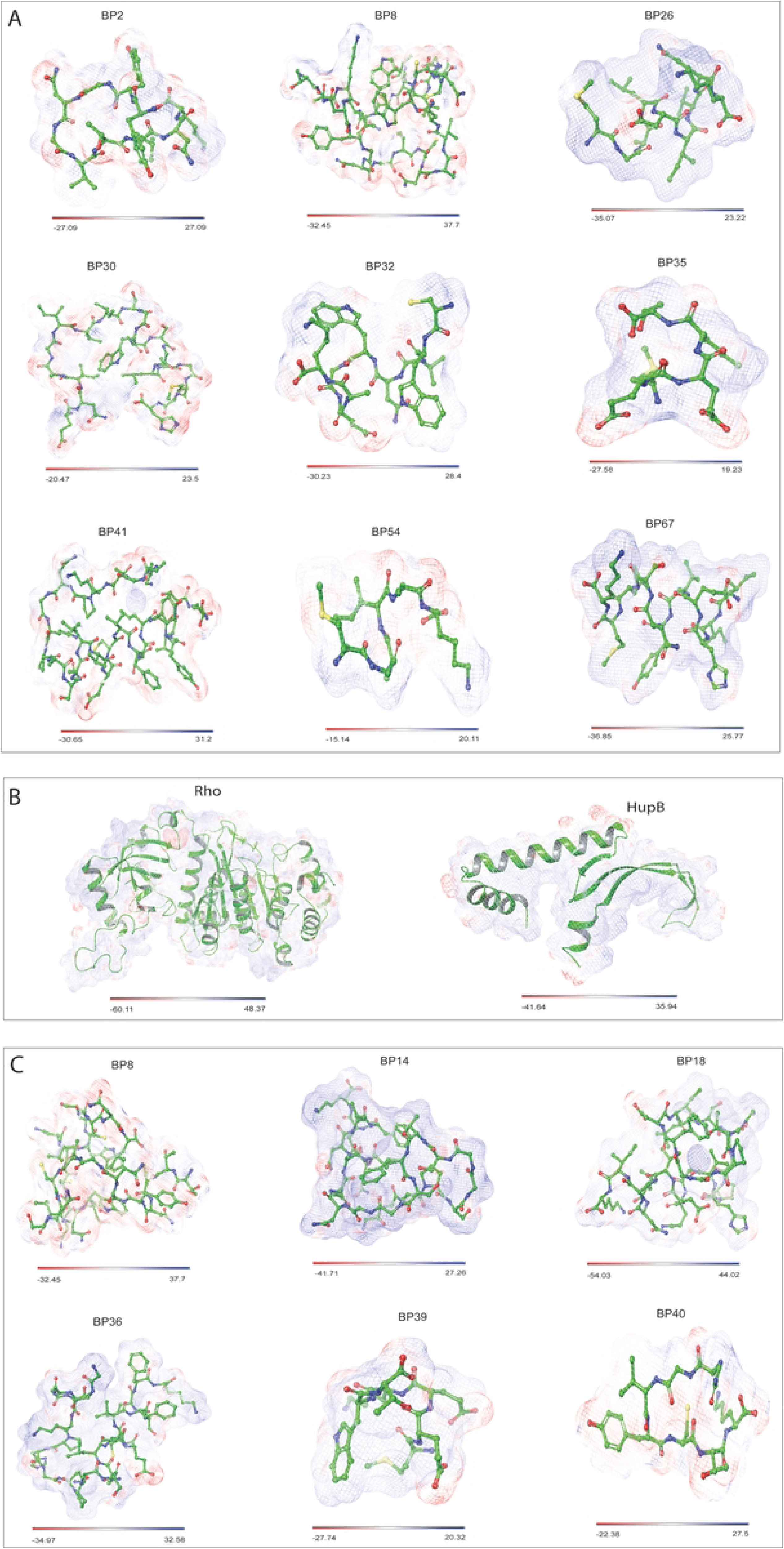
Electrostatic potential surface (EPS) of bacteriocin peptides (BPs) interacting with Rho and HupB proteins. (A) BPs interacting with Rho protein, (B) EPS of Rho and HupB proteins, and (C) BPs interacting with HupB protein. The EPS distribution is color-coded to show charge variations: red for negative, white for neutral, and blue for positive regions. This visualization highlights electrostatic interactions and provides insights into the binding and interaction sites between the molecules. All the EPS values are expressed as atomic unit (a.u.).

### Molecular dynamic simulation identifies potential inhibitory BPs against Rho and HupB proteins

The EPS analysis provided us with a detailed insights into the surface characteristics of both the BPs and their target proteins. Subsequently, molecular screening confirmed initial interactions between the BPs and the Rho and HupB proteins. To further understand these interactions and evaluate the thermodynamic stability of the BP-protein complexes, we conducted 200 ns molecular dynamics simulations for each complex. The simulation trajectories of the BP2-Rho complex revealed its overall stability over a 200 ns simulation period. The RMSD curve initially fluctuated up to 4Å, decreased to 3Å, and eventually reached 5Å (**Fig. 7A**) by the end of the trajectory, with some instability observed towards the end (**S1 File**). The BP8-Rho complex was the most stable, with RMSD fluctuations staying around 4Å and showing no significant deviations, confirming its thermodynamic stability (**Fig. 7B** and **S1 File**). For BP26-Rho, the RMSD initially peaked at 5Å but stabilized at 2.8Å after 80 ns, indicating improved stability as the simulation progressed (**Fig. 7C** and **S1 File**). The BP30-Rho dynamics exhibited RMSD fluctuations between 4Å and 5Å, indicating variability in the stability of the BP30-Rho complex (**Fig. 7D**). The animation further illustrated these fluctuations, highlighting significant instability, as the peptide displayed noticeable changes in its binding position over time (**S1 File**). BP32 demonstrated strong binding to the Rho protein, maintaining its position at the initial binding site with RMSD values consistently within the 4Å range, suggesting a stable interaction (**Fig. 7E** and **S1 File**). BP35 exhibited a stable interaction with the Rho protein, with RMSD values at 3Å for up to 150 ns before moving away from the initial binding site (**Fig. 7F** and **S1 File**). The RMSD plot for the BP41-Rho complex showed fluctuations reaching up to 5Å. The animation further confirmed these fluctuations, revealing some instability as BP41 interacted with the Rho protein (**Fig. 7G** and **S1 File**). BP54 maintained stability up to 100 ns, after which a slight displacement from the initial binding site was observed (**Fig. 7H** and **S1 File**). The BP67-Rho complex, like BP8-Rho, was among the most stable, as evidenced by the RMSD analysis and trajectory animations (**Fig. 7I** and **S1 File**). Overall, BP8 and BP67 were the most robust and stable complexes among the BP-Rho interactions studied. The initial molecular screening identified six BPs that effectively interacted with the HupB protein. Among these, BP8 showed the most stable interaction, with RMSD fluctuations up to 3.5 Å, indicating complex stability (**Fig. 8A**). This stability was corroborated by trajectory animations, which demonstrated consistent interactions between BP8 and HupB (**S2 File**). In contrast, the BP14-HupB complex displayed less stability, with higher RMSD fluctuations and inconsistent interactions in the animations (**Fig. 8B** and **S2 File**). The BP18-HupB (**Fig. 8C**) and BP36-HupB (**Fig. 8D**) complexes also showed significant RMSD fluctuations and lack of observable interactions (**S2 File**). Notably, BP39 and BP40 exhibited strong and stable interactions with HupB (**Figs. 8E-F**), with their 200 ns simulation movie revealing robust binding within the HupB binding pocket (**S2 File**).

**Fig. 7.**
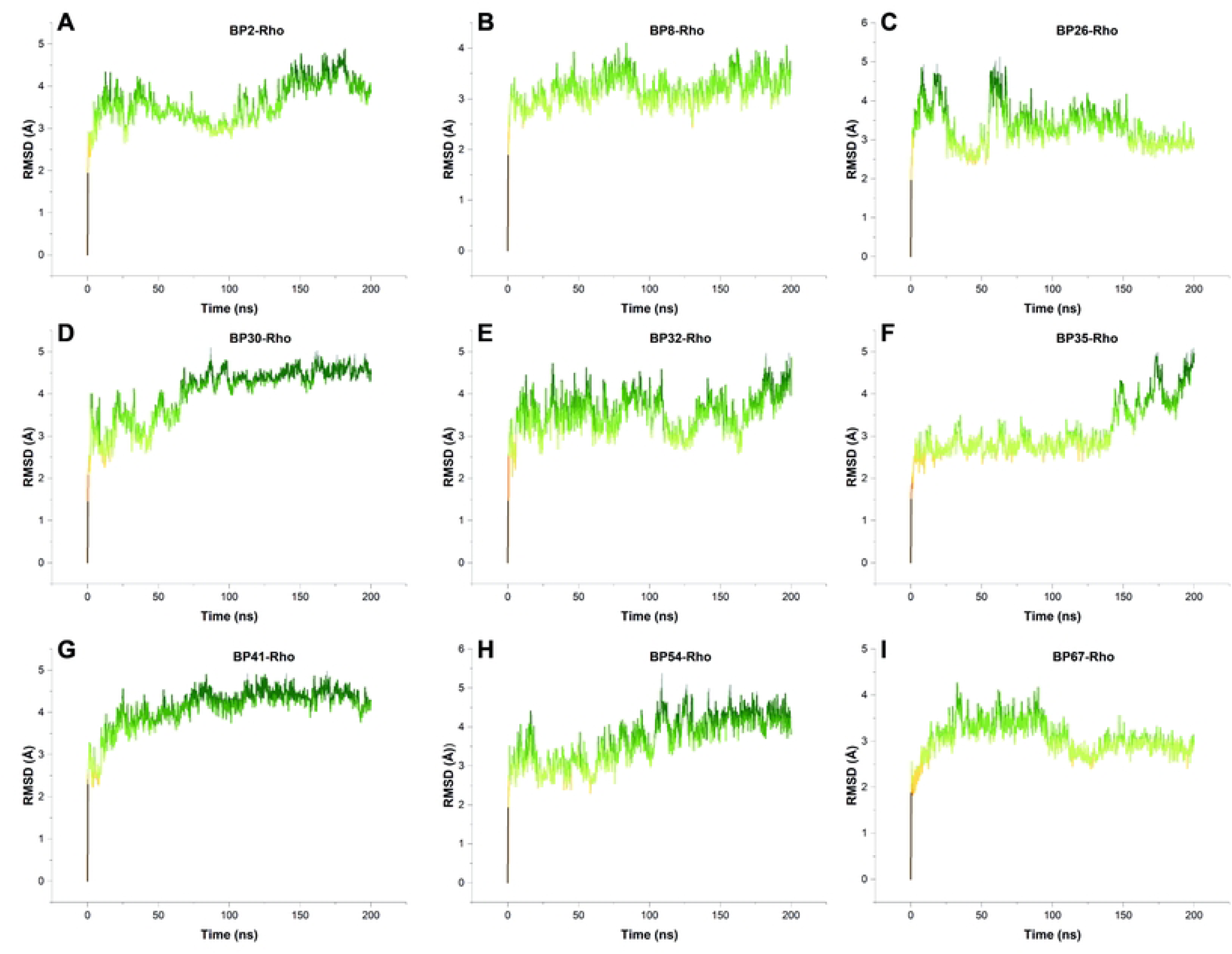
RMSD curves for BP-Rho complexes, showing structural stability throughout the dynamic trajectories. (A) BP2-Rho complex, (B) BP8-Rho complex, (C) BP26-Rho complex, (D) BP30-Rho complex, (E) BP32-Rho complex, (F) BP35-Rho complex, (G) BP41-Rho complex, (H) BP54-Rho complex, and (I) BP67-Rho complex. RMSD values are color-coded: light green indicates lower RMSD (more stable), and dark green denotes higher RMSD (slightly unstable).

**Fig. 8.**
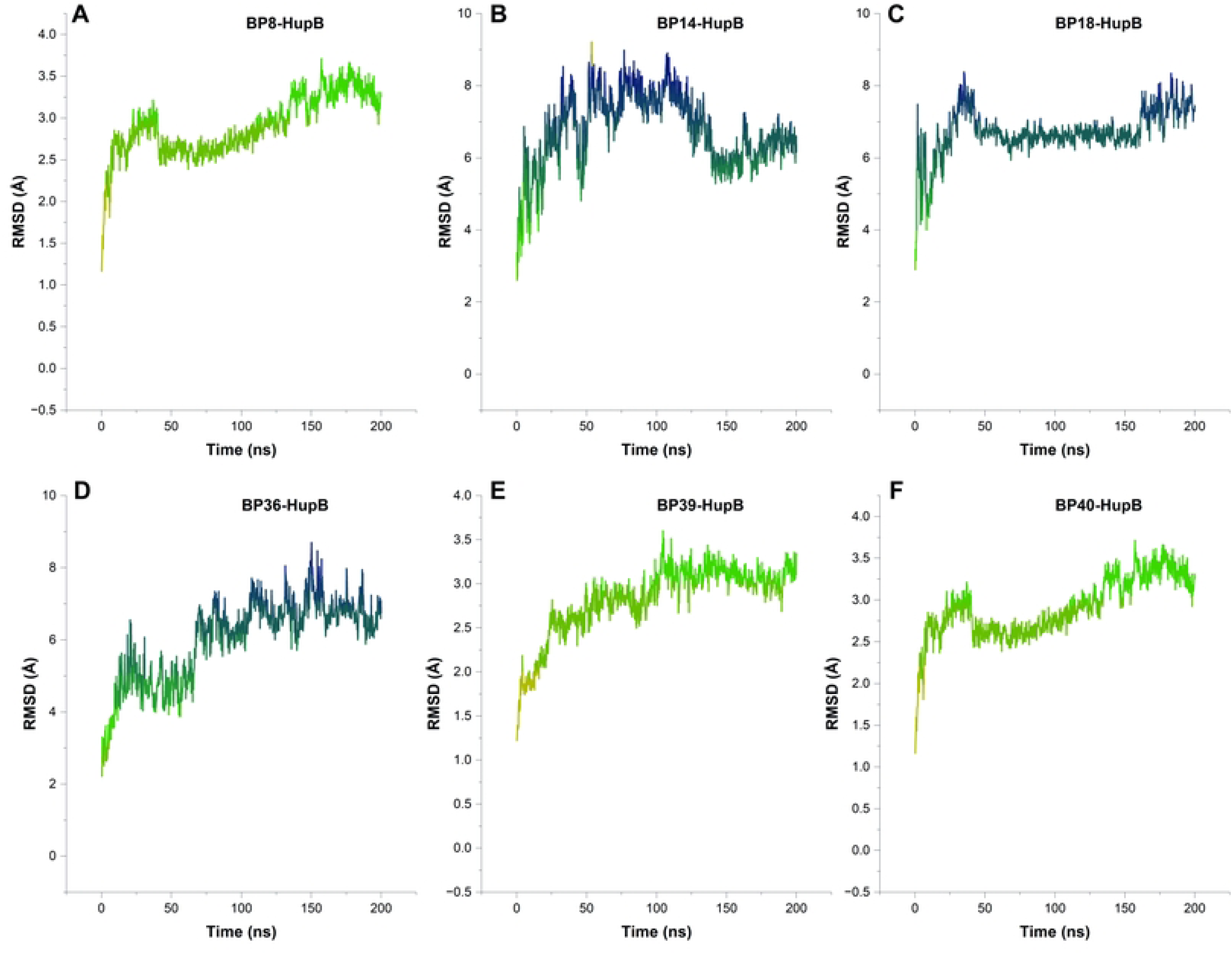
RMSD curves for BP-Rho complexes, showing structural stability throughout the dynamic trajectories. (A) BP8-HupB complex, (B) BP14-HupB complex, (C) BP18-HupB complex, (D) BP36-HupB complex, (E) BP39-HupB complex, and (F) BP40-HupB complex. RMSD values are color-coded: green indicates lower RMSD (stable), and increased density of greenish blue curves denote higher RMSD (unstable).

### Two BPs (BP8 and BP32) emerge as feasible inhibitors against Rho protein of mastitis pathogens

Each chain within the hexameric Rho protein interacts with central RNA, with ADP molecules bound in their respective ADP-binding clefts. For our analysis, we isolated the A chain of the Rho protein from the hexamer along with its associated RNA and ADP. Initial screenings indicated that the RNA maintained stable interactions with the Rho protein throughout the dynamic environment (**Fig. 9A**). Likewise, the ADP molecule displayed energetically favorable interactions within the ADP-binding pocket of Rho, maintaining stability throughout the simulation environment. Key residues implicated in ADP binding comprised LYS181, ALA182, GLY183, MSE186, ARG347, ALA350, GLU351, ARG353, VAL354, and PHE355 (**Fig. 9B**). Notably, in the presence of BP8, RNA ceased to interact with the Rho protein. Instead, BP8 displaced the RNA and formed interactions with Rho that mirrored the similar interactions depicted in **Fig. 4B**. For BP32, it displaced the ADP molecule, thereby preventing ADP from interacting with the Rho protein. This interaction resembled the previously reported Rho-BP32 interaction (**Fig. 4E**).

**Fig. 9.**
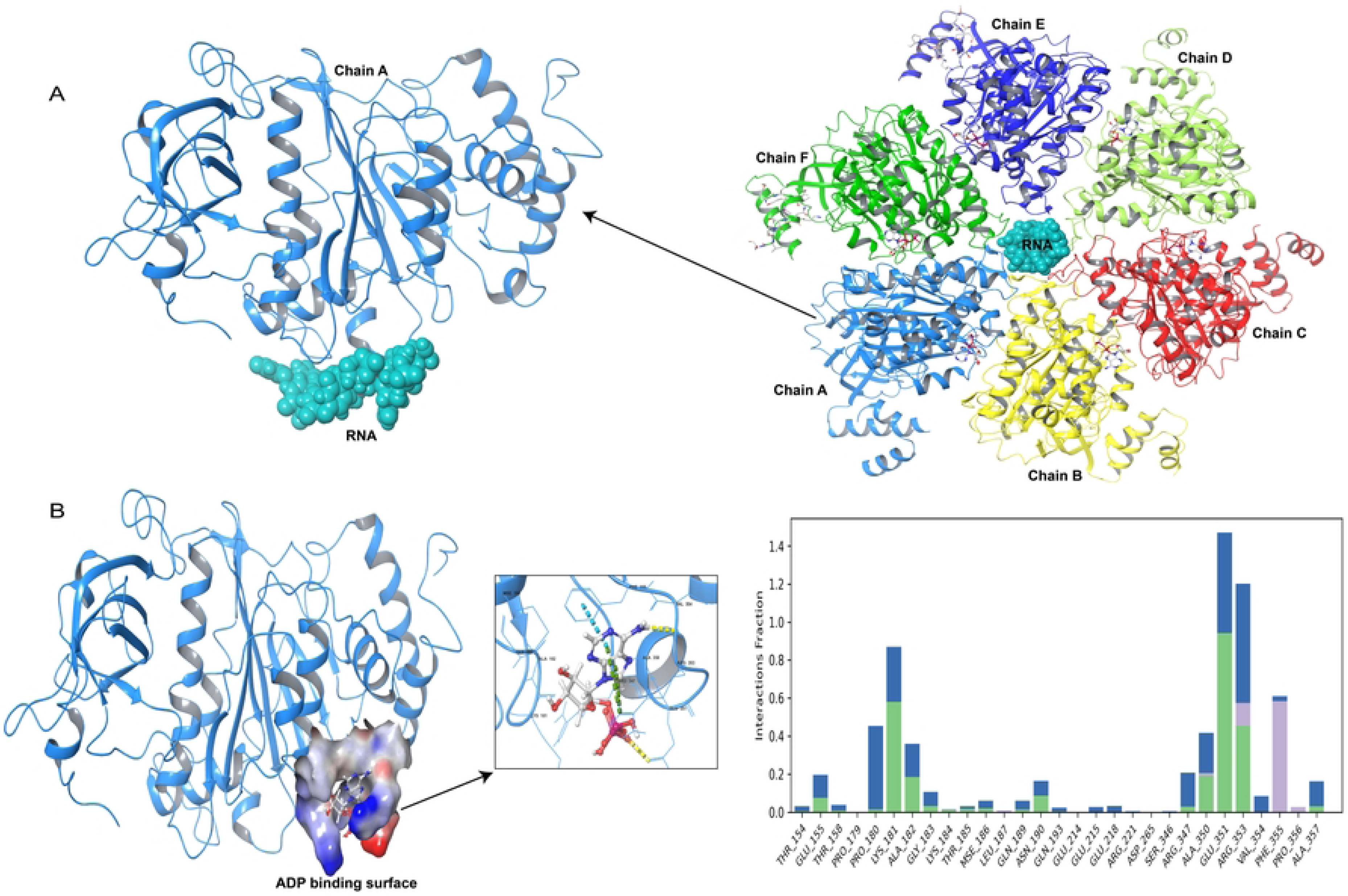
Interaction of hexameric Rho protein with RNA and ADP molecules. (A) The hexameric Rho protein consists of six chains, each interacting with a central RNA molecule, and each chain contains a distinct ADP-binding cleft. (B) The ADP molecule binds to the ADP-binding cleft through strong interactions with key amino acids, including LYS181, ALA182, GLY183, MSE186, ARG347, ALA350, GLU351, ARG353, VAL354, and PHE355.

## Discussion

Mastitis causes substantial economic losses in the dairy industry, with global annual losses estimated between $19.7 billion and $32 billion USD (3). The treatment of mastitis primarily depends on antibiotics. However, the increasing problem of AMR is a global concern, leading to a rapid decline in the availability of effective treatments against pathogenic bacteria (76, 77). While antibiotics have traditionally been used to treat infections (77), there is renewed hope in bacteriocins, natural antimicrobial agents that have already shown considerable effectiveness (78, 79). In this study, we introduced an innovative strategy to identify potent therapeutic agents targeting clinical mastitis-causing pathogens, uncovering highly effective antimicrobial peptides derived from bacteriocins through the integration of cutting-edge computational biology techniques and in-depth *in-silico* analyses. In this comprehensive investigation, we identified two conserved therapeutic targets (Rho and HupB), shared across 16 CM-causing pathogens in dairy cows (**S1 Table**). Furthermore, we predicted highly promising bacteriocin-derived inhibitory peptides (BPs) against these targets, with strong potential to target and disrupt these critical pathogen mechanisms, offering a novel approach for effective mastitis treatment. Utilizing a core genomic approach, we identified two essential proteins: Rho, a key transcription termination factor in bacteria (31, 32), and HupB, a nucleoid-associated protein critical for pathogen survival (34). The Rho and HupB proteins are predicted to be crucial for the survival of bacteria (34, 80), including CM-pathogens. They play essential roles in transcription and replication, with Rho facilitating transcription termination to regulate gene expression vital for bacterial growth (30), and HupB maintaining DNA structure and compaction necessary for effective replication (34). Disruption of these proteins could severely compromise the viability and pathogenicity of CM-causing bacteria, as evidenced by gene enrichment analysis (**Fig. 2**). Despite their crucial roles, the core proteins Rho and HupB have not yet been recognized as potential therapeutic targets for mastitis-causing pathogens. Our *in-silico* study is the first to predict their essentiality and virulence in CM-associated pathogens, utilizing established methods (81, 82) for identifying therapeutic targets. Structural comparisons of the 16 Rho and HupB proteins showed that their functional regions were nearly identical (**Figs. 3A** and **3D**). Since all these proteins were reported to perform the similar biological function, we classified them as isoforms with identical biological roles (83). Given their nearly identical structures, a single BP could potentially inhibit the function of all 16 pathogens by targeting their Rho and HupB proteins. The observed structural similarity between these proteins, coupled with the availability of their crystallographic structures, lends strong support to our findings and enhances the potential for their consideration as therapeutic targets.

Bacteriocins, known for their ability to inhibit bacterial growth (39), were initially selected and then fragmented into smaller BPs to identify their most active components. After screening 70 BPs against the two core proteins, Rho and HupB, we found that 14 BPs were effective in targeting these proteins. Among these, BP8 was particularly effective, demonstrating significant inhibitory activity against both Rho and HupB proteins. The findings of this study highlights that during the interactions between BP8 and the target proteins, both covalent and non-covalent bonds were prominently involved, underscoring the complex nature of these interactions and their potential significance in therapeutic applications (**Figs. 4** and **5**). However, the initial biochemical analysis of BPs showed varying results regarding the biochemical properties of the 14 BPs. The stability of the BP-receptor complexes remained unclear due to these initial interactions and the mixed biochemical data. Therefore, we employed a well-established MDS methods (71, 75) to accurately confirm the most effective BP compounds against Rho and HupB. Our study highlights BP8, BP32 and BP67 as notably stable complexes with Rho, maintaining low RMSD values (∼4Å) and minimal deviations, indicating significant thermodynamic stability. BP8 further demonstrates strong and reliable binding affinity with HupB, suggesting dual efficacy against pathogenic bacteria (**Figs. 7** and **8**). These findings underscore the potential of BP8 as a versatile therapeutic agent targeting multiple core proteins with stable interactions, supported by electrostatic complementarity (**S1** and **S2 Files**).

One of the important findings of this *in-silico* study is the identification of two BPs namely BP8 and BP32, which found to disrupt Rho protein function. The proper functioning of the Rho protein relies on its interactions with RNA and ADP molecules (84). We observed that the binding of BP8 to Rho prevents its interaction with RNA (**Figs. 4B** and **9A**). Similarly, BP32 inhibited ADP binding by occupying the ADP-binding site on the protein (**Figs. 4E** and **9B**). These findings underscore the potential of BP8 and BP32 as novel inhibitors targeting the Rho protein, a critical factor in bacterial pathogenesis (85). Regarding the HupB protein, while its precise role as a NAP remains incompletely understood, its involvement in the DNA replication process is well established (33, 86). Our analysis revealed strong binding interactions of three BPs (BP8, BP39, and BP40) with HupB, suggesting that their binding could effectively disrupt or terminate the activity of HupB. Specifically, their interaction with positively charged and polar residues such as LYS and THR (**Fig. 5**) could interfere with DNA binding of HupB, as these residues typically interact with the negatively charged phosphate groups of DNA (33). Furthermore, the small size of HupB increases the likelihood that these binding partners could effectively inhibit its function. *In-silico* studies of BPs against major mastitis pathogens are significant because they allow for the identification and optimization of effective peptides in a computational environment (38, 87), reducing the need for extensive *in-vivo* testing. These studies can predict BP’s efficacy, stability, and interactions with target pathogens, accelerating the development of targeted and sustainable treatments for mastitis (38, 87). With the increasing challenge of AMR and the known environmental impact of broad-spectrum antibiotics, there is a clear need for alternative treatments. We predicted Rho and HupB as novel therapeutic targets essential for bacterial survival (30, 35). Our findings indicated that BP8 and BP32 could effectively inhibit the activity of the Rho protein. Additionally, we hypothesized that BP8, BP39, and BP40 might disrupt the interaction of HupB in the bacterial DNA replication process. These peptides are of interest in various fields, including food preservation, probiotics, and medical applications (40, 78), due to their potential as natural alternatives to conventional antibiotics. While our approach focuses on treating bovine mastitis, it has potential for broader application against other pathogens of humans and animals.

## Conclusion

Mastitis remains a major challenge for the global dairy industry, particularly due to the escalating problem of AMR among the diverse pathogens implicated in this disease. With the growing demand for effective and environmentally sustainable antimicrobials, bacteriocins emerge as a compelling avenue for exploration. Utilizing a robust *in-silico* computational approach, we successfully identified six BPs with potent activity against the core proteins Rho and HupB across 16 key CM-associated pathogens. Among these BPs, BP8 and BP32 are predicted to disrupt Rho protein function by inhibiting RNA and ADP binding, while BP8, BP39, and BP40 prevent HupB from binding to DNA, making them potential inhibitors of mastitis pathogens. Thus, BP8 exhibited dual efficacy against both Rho and HupB, suggesting a multifaceted mechanism of action that enhances its therapeutic potential. These findings not only underscore the promise of BPs as a novel class of targeted antimicrobials but also introduce an innovative therapeutic strategy that could redefine mastitis management. By leveraging BPs, we may address the limitations of conventional antibiotics, thereby reducing reliance on traditional antimicrobials and mitigating the spread of resistance. However, while these *in-silico* findings are promising, they necessitate rigorous *in-vitro* and *in-vivo* validation to confirm efficacy, safety, and practical applicability in real-world settings. Further research in this area could pave the way for the development of next-generation therapeutics using BPs, offering a sustainable and effective solution to combat mastitis and other antibiotic-resistant bacterial infections in the future.

## Supplementary Information

**S1 Table**. Detailed information on the sixteen (N = 16) whole genome sequences of mastitis causing pathogens.

**S2 Table.** The average BLASTp results for identified ten essential core proteins (N = 10). These proteins were present in all of the 16 clinical mastitis-causing pathogens.

**S3 Table**. Protein information (peptide ID, sequence, number of amino acids, molecular wight, isoelectric point, net charge at pH, hydrophobicity, bioactivity etc.).

**S1 File.** Animation movies of top interacting BPs with Rho protein.

**S2 File.** Animation movies of top interacting BPs with HupB protein.

## Data Availability

The genomes sequenced, submitted to NCBI GenBank, and analyzed in this study are listed in S1 Table. Additional supporting data for this article can be found in S2 and S3 Tables, along with animated movie files (S1 and S2 Files) accessible at https://doi.org/10.6084/m9.figshare.27050914.

## Ethics Statement

This study was reviewed and approved by the Animal Research Ethics Committee (AREC) of the Bangabandhu Sheikh Mujibur Rahman Agricultural University, Bangladesh (Reference number: FVMAS/AREC/2023/6679, Date: 16/01/2023).

## Author Contributions

SH and MNH conceptualized and designed the study; SH, MMR and FY performed experiment, acquired and analyzed data, interpreted results and drafted the original manuscript; MJ, YAH, TI and MNH critically reviewed and edited the manuscript; all authors read, edited and approved the manuscript.

## Acknowledgments

We sincerely thank the dairy farmers for providing samples from their dairy cows, which were crucial for bacterial isolation, identification, and genome sequencing. Your support greatly contributed to the success of this research.

